# The characteristics of CTCF binding sequences contribute to enhancer blocking activity

**DOI:** 10.1101/2023.09.06.556325

**Authors:** Felice H. Tsang, Rosa J. Stolper, Muhammad Hanifi, Lucy J. Cornell, Helena S. Francis, Benjamin Davies, Douglas R. Higgs, Mira T. Kassouf

**Affiliations:** Chinese Academy of Medical Sciences Oxford Institute, Nuffield Department of Medicine, University of Oxford, Old Road Campus, Oxford, OX3 7BN, UK; MRC Weatherall Institute of Molecular Medicine, Radcliffe Department of Medicine, University of Oxford, John Radcliffe Hospital, Oxford, OX3 9DS, UK; Wellcome Centre for Human Genetics, Nuffield Department of Medicine, University of Oxford, Old Road Campus, Oxford, OX3 7BN, UK

## Abstract

While the elements encoding enhancers and promoters have been relatively well studied, the full spectrum of insulator elements which bind the CCCTC binding factor (CTCF), is relatively poorly characterised. This is partly due to the genomic context of CTCF sites greatly influencing their roles and activity. Here we have developed an experimental system to determine the ability of consistently sized, individual CTCF elements to interpose between enhancers and promoters and thereby reduce gene expression during differentiation. Importantly, each element is tested in the identical location thereby minimising the effect of genomic context. We found no correlation between the ability of CTCF elements to block enhancer-promoter activity with the amount of CTCF or cohesin bound at the natural genomic sites of these elements; the degree of evolutionary conservation; or their resemblance to the consensus core sequences. Nevertheless, we have shown that the strongest enhancer-promoter blockers include a previously described bound element lying upstream of the CTCF core motif. In addition, we found other uncharacterised DNaseI footprints located close to the core motif that may affect function. We have developed an assay of CTCF sequences which will enable researchers to sub-classify CTCF elements in a uniform and unbiased way.

## INTRODUCTION

The regulation of gene expression is controlled by two general classes of *cis*-acting elements: enhancers and promoters. However, within each class there is considerable variation in activity. For example, individual promoters and enhancers can be considered to be strong or weak, and as widely active or cell-type restricted. These features can to some extent be predicted from their sequence composition, ability to bind specific transcription factors and epigenetic signatures (1). However, there may be overlap between these classes of element: to some extent all enhancers may act as promoters and some promoters can act as enhancers (2–4). The activities of enhancers and promoters are modified by a third class of fundamental *cis*-acting elements referred to as CCCTC binding factor (CTCF) elements which have a wide range of activities including: blocking interactions between enhancers and promoters (5–7); facilitating interactions between enhancers and promoters (8–11); acting as barriers between active and inactive chromatin (12–14); contributing directly to activity of enhancers and promoters; and playing a key role in the three dimensional structure of the genome thereby influencing the physical proximity of enhancers and promoters (15–17). The genome contains 20,000-50,000 CTCF binding sites, but other than binding CTCF (14,18,19), relatively little is known about how to classify individual elements and to assess their ability to perform each aspect of their many potential activities. Previous studies derived from large datasets have correlated the activities of CTCF elements with features such as the core CTCF sequences, its evolutionary conservation, the amount of CTCF and cohesin enriched at the CTCF sites, and the persistency of CTCF binding after CTCF depletion (18,20–24). However, the predictive values of these parameters have not been extensively tested on individual elements.

Assays of enhancers and promoters in terms of sequence, epigenetic modification and function are relatively well developed and standardised whereas assays of CTCF elements are less well characterised, in part because they have such diverse functions, and their role greatly depends on their genomic context. For example, for a CTCF element to block the activity of an enhancer on its cognate promoter it has to lie between such elements (5–7), whereas for CTCF elements to facilitate enhancer-promoter interactions they must lie within or close to the interacting sequences (8–10). We have recently investigated the roles of CTCF sites within and surrounding a ∼65kb sub-TAD containing the mouse alpha-globin cluster (Figure 1a) by investigating how deletion of individual sites and combinations of sites affect 3D structure of the sub-TAD and alpha globin expression (25,26). None of these sites appear to influence the levels of alpha globin expression but two sites (HS-38 and HS-39) insulate neighbouring genes (Mpg, Rhbd1, Il9r) from the very strong cluster of enhancers controlling alpha-globin expression (26).

**Figure 1.**
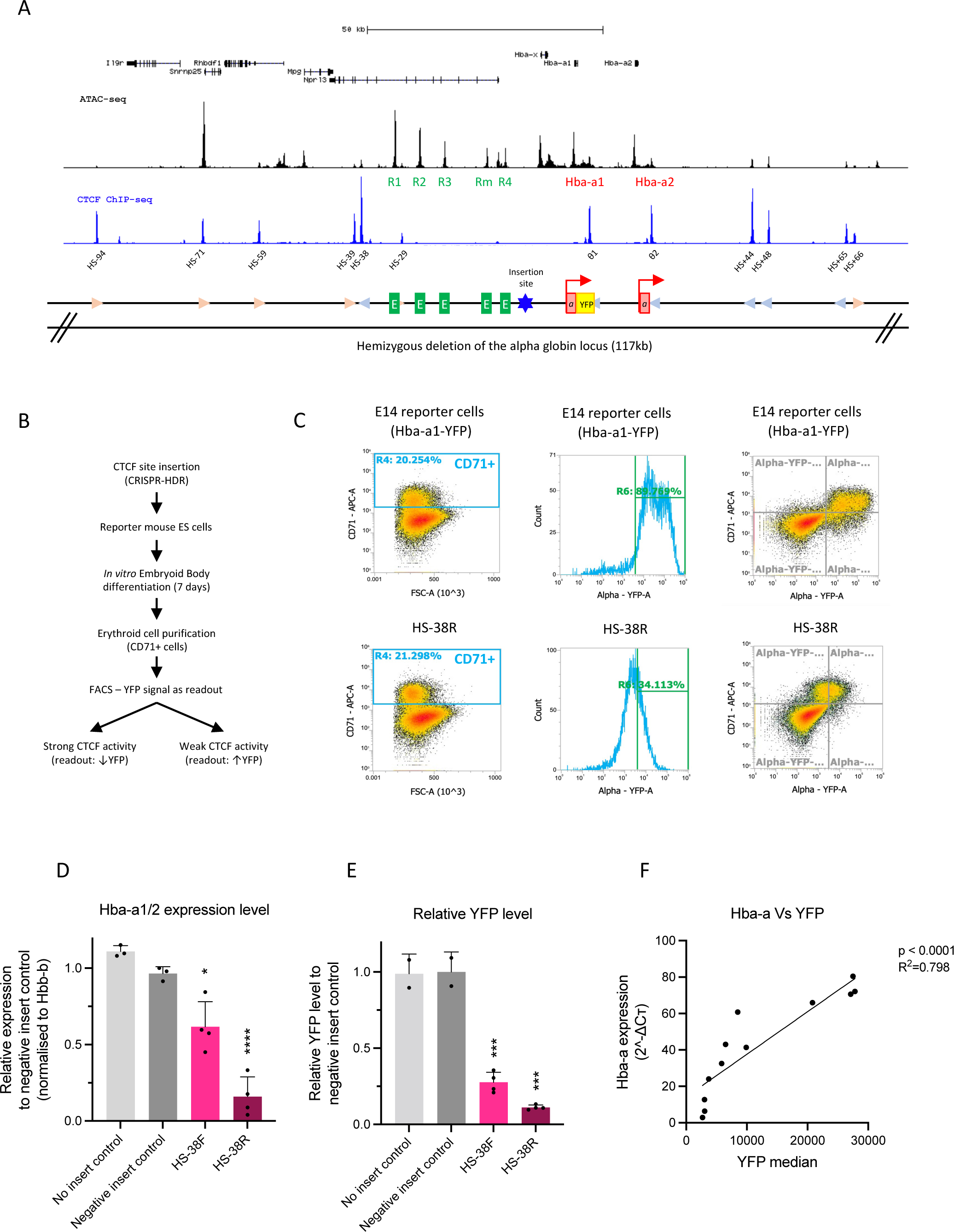
An experimental system to evaluate the capacity of individual CTCF elements to block enhancer-promoter interactions. (a) The CTCF site of interest was inserted between the enhancer clusters and the promoters at the mouse alpha-globin locus. An allele of the alpha-globin gene locus (117kb) was hemizygously deleted in the reporter mouse ES cells (mESCs) and the Hba-a1 gene was tagged with a YFP protein. (b) The CTCF site of interest was inserted into the hemizygous alpha-globin locus of the reporter mESCs by the CRISPR-HDR method. The modified reporter mESCs were differentiated into EBs and the cells were harvested for analysis. The YFP level of the CD71+ cell population in the EBs were determined by FACS analysis and interpreted as the readout of the CTCF ability to block enhancer-promoter interaction. (c) CD71+ cells isolated from the differentiated EBs exhibit high level of YFP signal as measured by FACS analysis. The upper and lower panels represent the FACS profile of the WT reporter mESCs and the cells with the inserted HS-38R site, respectively. (d) Insertion of the HS-38 sequence in both the forward and reverse orientation significantly represses the expression of the alpha-globin genes (*Hba-a1/2*) normalised to the *β*-globin (*Hbb-b*) when compared to the ‘no insert’ and ‘negative insert’ controls. (e) Insertion of the HS-38 sequence in both the forward and reverse orientation significantly represses the YFP level of the reporter cells when compared to the ‘no insert’ and ‘negative insert’ controls. (f) The expression of the alpha-globin genes is positively and significantly correlated to the level of the YFP in the samples (Linear regression, p<0.0001, R^2^=0.798).

In addition to this observation, we inserted HS-38 individually and in combination into a non-coding region of the sub-TAD lying between the alpha globin enhancers and promoters (insertion site in Figure 1a) (Stolper, Tsang et al.). In this position HS-38 acted to partially block the interaction between the enhancers and promoters and significantly reduced the level of alpha globin expression. Multiple insertions of HS-38 (2-4 copies) increasingly compromised enhancer-promoter interactions and further decreased alpha globin expression. Importantly, we demonstrated that the blocking activity of HS-38 and accumulation of cohesin at this site was greater when the N-terminus of bound CTCF was orientated towards the enhancers rather than towards the promoters, consistent with the bound CTCF stalling translocation of cohesin from the enhancers to mediate loop extrusion (Stolper, Tsang et al.). Some degree of enhancer-promoter blocking was also seen when the CTCF element was inserted in the opposite orientation (Stolper, Tsang et al.). Consistent with previous conclusions it is proposed that blocking the translocation of cohesin reduces the frequency of juxtaposition between enhancers and their cognate promoters within a TAD or sub-TAD (27,28).

Here, we have engineered the alpha globin cluster to enable a direct comparison and ranking of the extent to which individual CTCF elements can compromise enhancer-promoter activity and thereby reduce alpha globin expression during differentiation: just one of many activities of such sites. Here we have analysed a set of 18 CTCF elements whose roles have been previously determined by deletion from their natural genomic environment and/or analysed in other assays of CTCF elements.

## MATERIAL AND METHODS

### Construction of plasmids

To generate the CRISPR-Cas9 and chimeric guide RNA expression construct, the single guide-RNAs (sgRNAs) targeting the CTCF element insertion site were cloned into the pX335-U6-Chimeric_BB-CBh-hSpCas9n(D10A) (Addgene, 42335) vector as previously described (26). The pX335 vector was modified to contain a puromycin selection cassette. Sequences of the sgRNA are listed in table 1. Next, oligonucleotides corresponding to each CTCF element were cloned into the pROSA-TV2 vector (created by Prof. Benjamin Davies group) between the pair of BsaI sites using the Golden Gate Assembly Kit (NEB, E1601) and this generates the HDR donor template containing the inserted CTCF element. The pROSA-TV2 vector was designed to contain the 1.4Kb and 1.2Kb homology arms of the inserted site and a hygromycin selection cassette that is flanked by a pair of lox sites (loxP). Sequences of the homology arms are listed in the supplementary table 1 and the inserted CTCF element oligonucleotides are listed in the table 1.

**Table 1.**
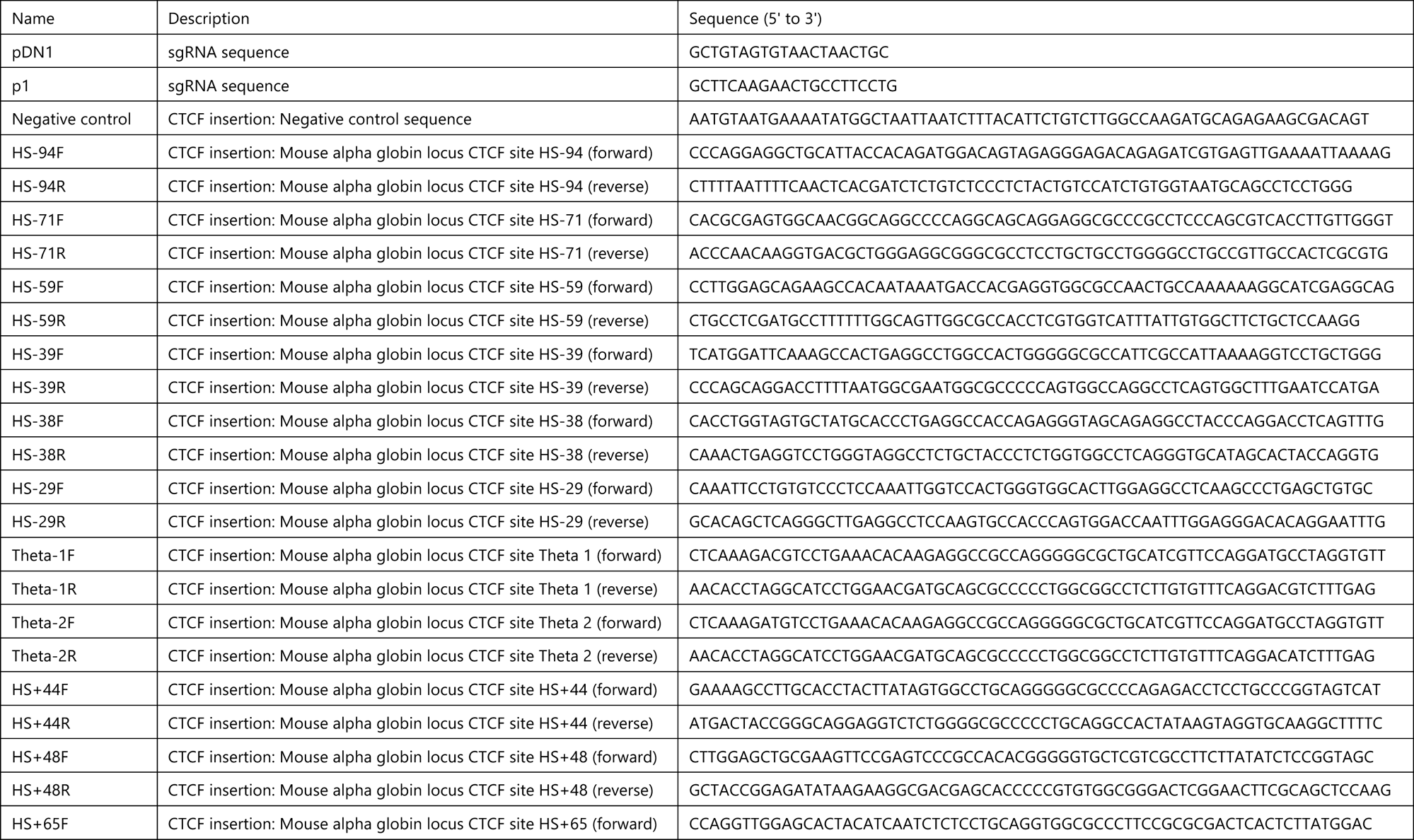
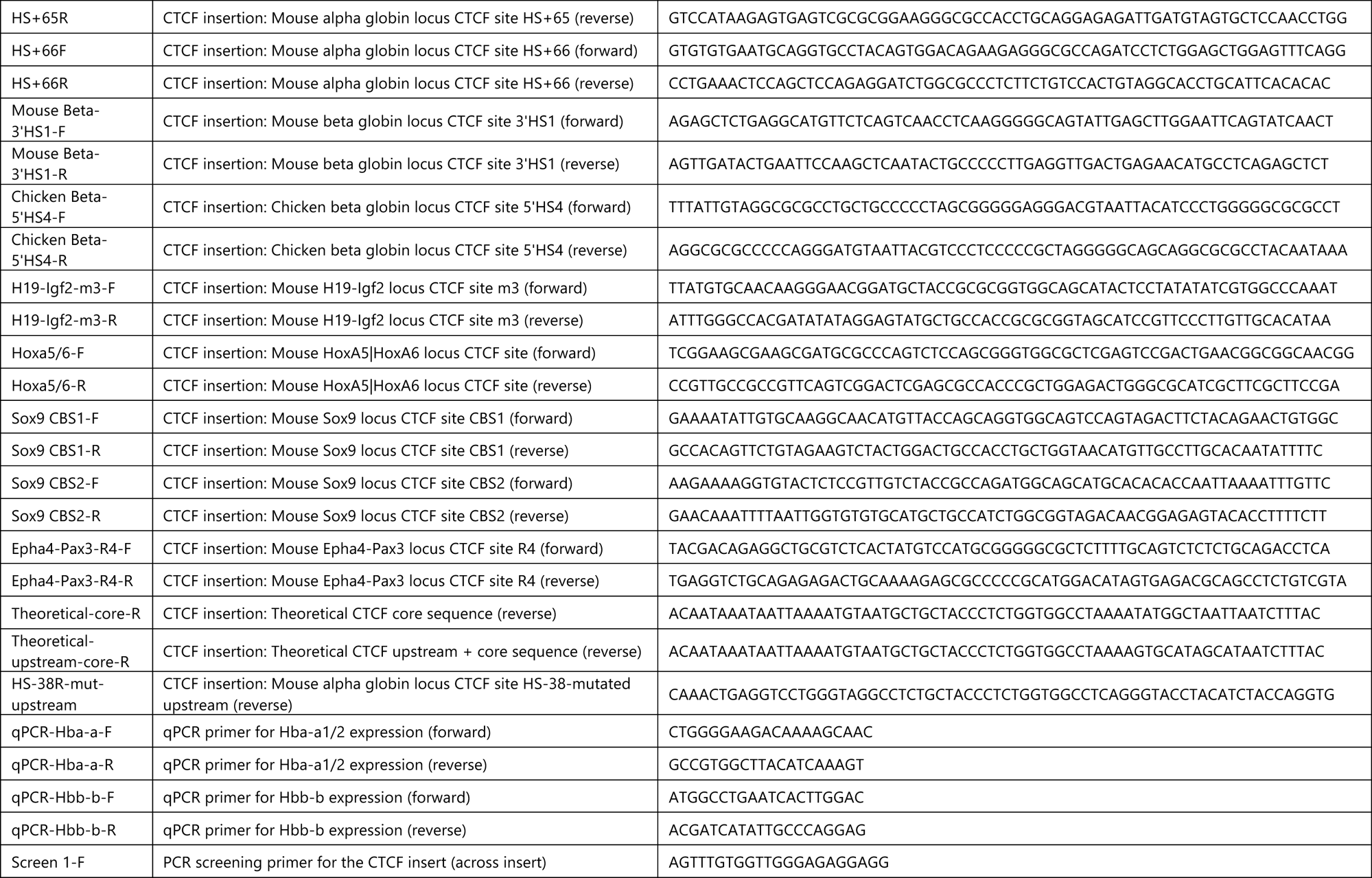
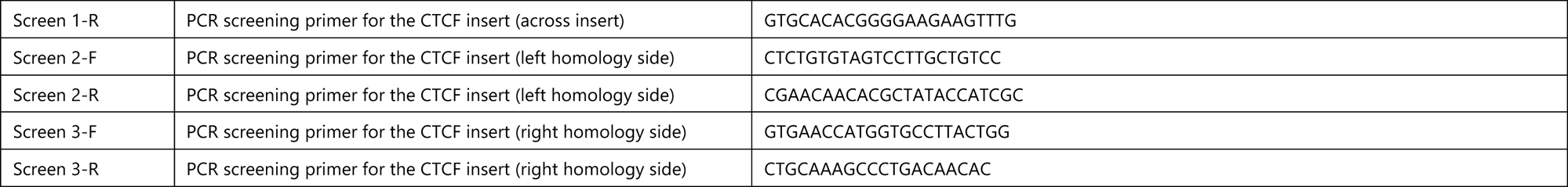
The sgRNAs, primers, and all the CTCF elements insertion sequences.

### Tissue culture, transfection and drug selection

The mouse embryonic stem cell line (mESC), E14TG2a was used for the CTCF activity reporter assays. To facilitate the genetic modifications and downstream analysis, one wild-type copy of the alpha-globin locus was hemizygously deleted, and a yellow fluorescent protein (mVenus) was introduced at the end of exon 3 of Hba-a1 gene (29). The mESCs were maintained in Glasgow’s Minimal Essential Medium (Gibco 21710-082) supplemented with 10% heat-inactivated foetal bovine serum (Gibco, 10270-106) 1mM Sodium pyruvate (Gibco 11360-039), 2mM L-Gluatamine (Gibco, 25030-024), 1x MEM (NE) AA (Gibco 11140-035), 0.1mM beta-mercaptoethanol (Gibco, 31350-010), and 1U/μl leukaemia inhibiting factors (Cell guidance system, LIF, GFM200) at 37°C humidified chamber with 5% CO_2_. Genetically modified mESC models with inserted CTCF elements were generated using the double nickase CRISPR-mediated homology-directed repair (HDR) strategy. The mESCs were co-transfected with the 1.66 μg of the two sgRNA nickase-Cas9 plasmids and 0.83 μg of the HDR plasmid using the Lipofectamine LTX and Plus Reagent (Invitrogen, 15338-100) according to manufacturer’s instructions. Transfected cells underwent 1 μg/ml puromycin selection for 2 days and a subsequent 250 μg/ml hygromycin selection for 6 days. The positively selected cells were then transfected with 2.5 μg pCre-Pac vector using the Lipofectamine LTX and Plus Reagent (Invitrogen, 15338-100) which facilitated removal of the hygromycin-resistance cassette.

### Genotyping

Genetically modified mESCs that survived the antibiotic selections and Cre-recombinase vector transfection were single cell sorted in 96-well plate and grown as clonal colonies. Genomic DNA was extracted from individual colonies and were subjected to 3 sets of PCR screening (across the insert, left homology side spanning, and right homology side spanning) for the successful insertion of the CTCF elements to the target site and to eliminate possible concatemer integrations (Thermo Scientific, DreamTaq Green PCR Master Mix, K1081) (Supp. figure 1b). Genotypes of positive clones were confirmed by Sanger sequencing. Sequences of the PCR screening primers are listed in table 1.

### *In vitro* Embryoid body (EB) differentiation

EB differentiation of mESCs were based on previously published protocol (29). In brief, mESCs were induced by passage into IMDM base media (Iscove’s modified Dulbecco’s medium (IMDM, Gibco, 31980030), supplemented with 15% heat-inactivated FBS, 1.4×10^−4^ M monothioglycerol (Sigma-Aldrich, MTG, M6145), 50 U/ml penicillin-streptomycin (Gibco, 15140122) and 1000 U/ml LIF) at 48 hours prior to differentiation. During primary plating, mESCs were trypsinized and plated in the EB differentiation medium (IMDM medium supplemented with 15% heat-inactivated FBS, 5% protein-free hybridoma medium (PFHM-II, Gibco, 12040-077), 2 mM L-glutamine, 50 μg/ml L-ascorbic acid (Sigma Aldrich, A4544), 3×10^−4^ M MTG and 300 μg/ml human transferrin (Roche, 10652202001)) in a triple vent petri 10 cm dishes (Thermo fisher, 101VR20) seeded at 1×10^4^ cells for up to seven days. The dishes were gently shaken daily to disrupt any potential attachment to the dish. At day 7, EBs were harvested and disaggregated in 0.25% trypsin (Gibco, 25200-056) for 3 minutes at 37°C and neutralized with FBS. The disaggregated EBs were labelled with anti-mouse CD71-FITC antibody (eBioscience, 11-0711-85) (1:200 in staining buffer), washed, and further incubated with the MACS anti-FITC separation microbeads (Miltenyi, 130-048-701, 10 μl per 10^7^ cells). The bead labelled CD71+ cells were then isolated by magnetic column separation (Miltenyi, LS Column, 130-042-401), according to the manufacturer’s instructions.

### Flow cytometry

Differentiated EB cells were desegregated and stained with the anti-mouse CD71-APC antibody (eBioscience, 11-0711-80) (1:8000 in staining buffer) and Hoechst (Invitrogen, 33258) (1:10000 in staining buffer) for 30 minutes at 4°C in dark. The level of YFP signal in the live CD71 positive cells was measured and used as a proxy for the level of the alpha-globin gene expression in cell line models. Stained cells were analysed using the Attune NxT Flow cytometer (Thermo fisher) and the Attune NxT software package.

### RNA expression analysis

Total RNA was isolated from 1×10^7^ EB-derived CD71+ cells by using TRI reagent (Sigma, T9424). RNA extraction and the in-column DNaseI digestion were performed by using the Direct-zol RNA miniprep kit, according to manufacturer’s protocol (Zymo Research, R2050). The quantity and quality of the RNA are accessed by the Qubit RNA BR Assay (Invitrogen, Q10211) and the RNA ScreenTape (Agilent Technologies, 5067-5576), respectively. 1μg of the total RNA was used for synthesizing cDNA using the Superscript III first-strand synthesis SuperMix (Invitrogen, 11752-050) according to manufacturer’s protocol, a no reverse transcriptase control with included to track potential genomic DNA contamination in the sample. Quantitative real-time PCR was performed in triplicate on the 5x diluted cDNA using the fast SYBR green master mix (Thermo Fisher, 4385612) to assess the relative changes in gene expression. Primers used in the qPCR reaction is listed in table 1. The ΔΔCt method was used quantify the RNA abundance of *Hba-a* relative to *Hbb-b*.

### CTCF core binding motif occurrence

The occurrence of the CTCF core binding motif was determined by the “*Find Individual Motif Occurrences*” FIMO (version5.4.1) program in the MEME suite (30). The CTCF core motif position weight matrices was based on the HOCOMOCO database (CTCF.MOUSE.H11MO.0.A) (31). The p-value generated by FIMO analysis represents the statistical significance of the occurrence of the CTCF core motif within a sequence. When multiple CTCF core motifs were detected within the sequence, only the best match was retained.

### Conservation of the CTCF element

The degree of evolutionary conservation of the 68bp inserted CTCF sequence was indicated by PhastCons conservation score. PhastCons is a hidden Markov model-based method that estimates the probability that each nucleotide belongs to a conserved element based on multiple alignment in 30 vertebrate species (32).

### Enrichment level of CTCF and Rad21

The enrichment level of CTCF protein and RAD21 protein on dividual CTCF binding sites were determined by re-analyzing the previously published CTCF and RAD21 ChIP-sequencing data in mESCs and erythroid cells with the Lanceotron peak calling framework (LanceOtron.molbiol.ox.ac.uk) (33). Peak calling was performed using the bigwig files of the ChIP-seq dataset as the input, and the enrichment level of CTCF or RAD21 of individual ChIP-seq peak is calculated as the peak statistics (width, height, area) in Lanceotron.

### Chromatin immunoprecipitation sequencing

Chromatin immunoprecipitation sequencing (ChIP-seq) was performed using the ChIP Assay Kit according to the manufacturer’s instructions (Milliopore, 17-295). In brief, 5×10^6^ purified CD71 positive erythroid cells derived from two biological replicates were subjected to crosslinking with 1% formaldehyde for 10 minutes and was quenched with 125mM of glycine for 5 minutes. Chromatin fragmentation was performed using the Bioruptor Pico Sonicator (Diagenode) with 6 cycles of 30s ON and 30s OFF at 4°C to obtain an average fragment size of 200-500 bp. The fragmented chromatin was immunoprecipitated with 10μl of Rabbit Anti-CTCF antibody (Millipore, 07-729) at 4°C overnight on a rotator. The mix was incubated with Protein A agarose beads for 1 hour at 4°C on a rotator prior to the washing steps and the elution. DNA sequencing libraries were prepared with the NEBNext Ultra II DNA library prep kit according to the manufacturer’s instructions (New England Biolabs, E7645S) and sequenced on the Illumina NextSeq platform with the 75-cycle paired end kit (NextSeq 500/550 High Output Kit).

ChIP-seq data was analysed using an in-house pipeline. In brief, bowtie (version 1.2.3) was used to align the raw fastq files with the reference genome (an edited customed genome based on mm10 that include the CTCF insertion sequence). The remaining unaligned reads are trimmed by Trimgalore (version 0.6.5) and flashed by using FLASh (version 1.2.11) before re-aligned to the reference genome by bowtie again. Samtools (version 1.10) was used to filter, sort, fixmate, remove PCR duplicates and index the mapped reads. The bamCoverage function of Deeptools (version 2.2.2) was used to create the bigwig file with normalisation set to CPM and smooth length of 300bp.The bigwig files were visualized on the University of California Santa Cruz (UCSC) genome browser as individual tracks for comparison. All ChIP-seq experiments were performed at least in biological duplicate with similar results.

### Statistical analysis

Statistical analysis was carried out with Graphpad Prism (version 9). All the experiments were performed on three biological replicates (3 individual clones and 3 individual differentiations) with similar results, and standard deviation is shown for all measurements. Statistical analysis between control and target groups was performed using unpaired, two-tailed *t*-tests. Spearman correlation was performed for the FIMO p-value and CTCF enhancer-promoter blocking activity. Linear regression was performed to determine the relationship between the CTCF enhancer-promoter blocking activity and variables including the degree of conservation, levels of CTCF enrichment, and level of Rad21 enrichment. *P-*values are represented as ^NS^ P > 0.05; ^∗^ *P* < 0.05; ^∗∗^ *P* < 0.01; ^∗∗∗^ *P* < 0.001; ^∗∗∗∗^ *P* < 0.0001.

## RESULTS

### An experimental system to evaluate the capacity of individual CTCF elements to block enhancer-promoter interactions

Using the mouse alpha-globin locus, we developed an experimental system to quantify the strength and effectiveness of individual CTCF elements to block the interaction between enhancers and promoters during erythroid differentiation. The regulatory landscape of the mouse alpha-globin locus has been well-characterized and the protocol for erythroid differentiation is fully established (29). We have previously shown that the duplicated alpha-globin genes are regulated by a cluster of five enhancers (R1, R2, R3, Rm, and R4) located 14-37 Kb upstream of the *Hba-a1* gene. The alpha-globin enhancers and genes are located within a ∼65kb sub-TAD (topologically associating domain) that is largely delimited by an array of convergently oriented CTCF binding sites (26,34,35) (Figure 1a).

We have recently shown that insertion of an 83 bp fragment spanning a previously characterised CTCF binding element (HS-38) between the alpha-globin enhancers and the alpha-globin promoters significantly reduced the interaction between these elements and reduced the level of transcription from the alpha-globin genes (Stolper, Tsang et al.). This CTCF element acted in an orientation dependent manner, having a greater effect when the N-terminus of the bound CTCF protein was directed towards the upstream enhancers. Based on this study, we engineered the alpha-globin locus to allow insertion of any CTCF bound element into a landing pad placed in a non-coding region between the enhancers and promoters (insertion site in Figure 1a) to determine the ability of each independent element to reduce the interaction between the enhancers and their cognate promoters. To standardise the comparison of CTCF elements we analysed uniformly sized sequences (68 bp) containing the 20 bases core CTCF motif, 24 bases upstream and 24 bases downstream to include all known DNA binding surfaces of CTCF.

In undifferentiated mES cells, CTCF elements of interest were inserted into a landing pad (insertion site) between the alpha-globin enhancers and the alpha-globin promoters using CRISPR-mediated homology-directed repair (HDR) (Supp. figure 1a). To facilitate the CRISPR-HDR insertion and analysis of the engineered allele, we hemizygously deleted the wild-type copy of the alpha-globin locus (117 kb). In addition, we tagged the *Hba-a1* gene with a yellow fluorescent protein (YFP), so that the YFP signal could be used as a proxy for the level of alpha-globin gene expression (Figure 1a) (29). These engineered mES cells were subsequently differentiated into erythroid cells *in vitro* within embryoid bodies (EBs) (29) since the interaction between the alpha-globin enhancers and their cognate promoters only occurs in differentiating erythroid cells. The levels of YFP in differentiated CD71+ erythroid cells were quantified by flow cytometry as a readout of the degree to which any CTCF element acted as an enhancer blocker (Figure 1b and 1c): high levels of YFP indicate low levels of insulation and low levels of YFP are associated with high levels of insulation.

To validate this as a means of testing how individual CTCF elements could block enhancer-promoter interactions, we initially inserted the HS-38 CTCF binding site into the landing pad. This element had previously been shown to effectively reduce enhancer-promoter interactions at the alpha-globin locus (Stolper, Tsang et al.) where it normally insulates interaction between the strong alpha-globin enhancers and genes (Mpg, Rhbdf1 and Il9r) lying upstream of the cluster (26). Insertion of a 68bp element spanning the HS-38 CTCF element, in both the forward and reverse orientations exhibited significant reductions in alpha-globin gene expression when compared to the no insert control (the unmodified mESCs) and a negative insert control (insertion of the 68bp neutral sequence with no boundary activity) (Figure 1d). The reduction of alpha-globin expression was accurately reflected by the decreased level of YFP signal (Figure 1e) which is strongly and positively correlated with the level of alpha-globin RNA expression, assessed by RT-qPCR (p < 0.0001, R^2^=0.798) (Figure 1f). Consistent with the work of Stolper and Tsang et al., we observed that the ability of the CTCF element to block an interaction between the enhancers and promoters was greater when the HS-38 site was inserted in the reverse orientation with the N-terminus of bound CTCF orientated towards the upstream enhancers. This observation may reflect the directional tracking of the cohesin complex from the enhancers to the promoters on the alpha-globin locus as previously discussed (Stolper, Tsang et al). In summary, we have shown that this engineered cell line and the associated YFP reporter faithfully and accurately reflect the ability of an individual CTCF element to reduce the functional interaction between an enhancer and promoter as the cells undergo differentiation.

### Analysing previously characterised CTCF elements

To investigate whether this experimental model could distinguish the degree to which individual CTCF elements might differ in their ability to interfere with enhancer-promoter interactions, we analysed previously characterised elements. Our previous work deleting individual boundary elements in the alpha-globin locus showed that HS-38 acts as a strong boundary which normally delimits the activity of the alpha-globin enhancers on genes lying upstream of the cluster (26). Deletion of other CTCF elements in the alpha-globin sub-TAD had variable effects on its 3D structure. Importantly, deletion of individual sites and combinations of sites from their natural positions in the alpha-globin sub-TAD had no discernible effects on alpha-globin expression (25).

Insertion of individual CTCF elements of the alpha-globin locus in turn into the landing pad blocked the enhancer-promoter interaction to variable degrees (Figure 2a). Insertion of these boundary sequences in both forward and reverse directions showed similar trends (Figure 2b) but again with greater effect when the N-terminus of bound CTCF was directed towards the enhancers. These observations demonstrated that the experimental system established here can distinguish CTCF elements which display different degrees of insulation in an orientation-dependent manner.

**Figure 2.**
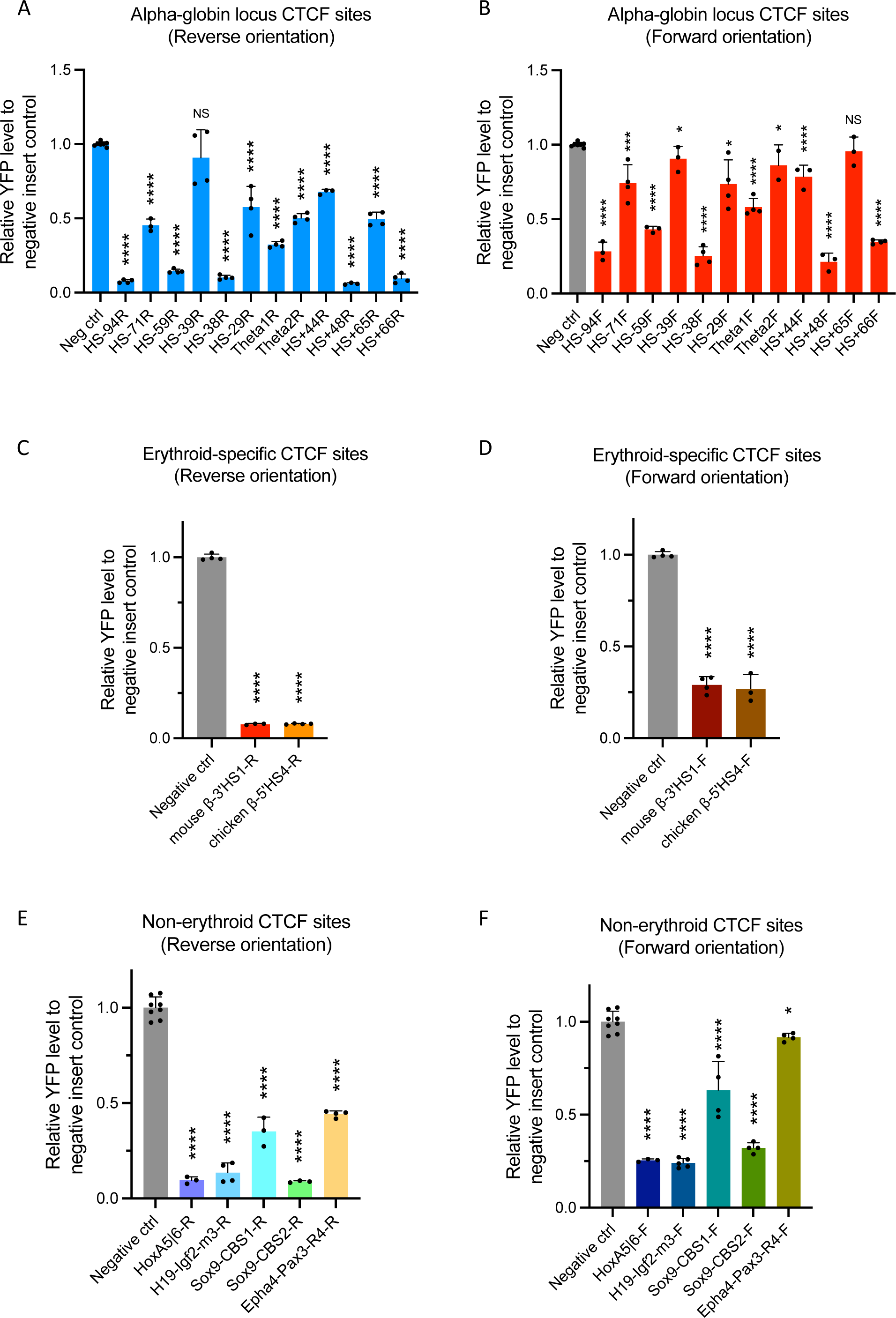
Testing previously characterised CTCF elements for their insulation activity. (a)&(b) Nearly all CTCF sites in the alpha-globin locus blocked the enhancer-promoter interaction to variable degrees in both orientations. (c)&(d) Both *β*-globin locus CTCF binding sites in mouse (3’HS1) and chicken (5’HS4) significantly blocked the enhancer-promoter interaction. (e)&(f) CTCF binding elements from other non-erythroid specific loci also blocked the enhancer-promoter interaction and reduced alpha-globin expression to variable degrees. (NS not significant; * p<0.05; ** p<0.01, *** p<0.001, **** p<0.0001)

Initial experiments were performed on CTCF elements derived from the mouse alpha-globin TAD. A more extensive set of CTCF elements was tested to ensure that the experimental system was suitable for analysing a broader range of elements. To this end, we first analysed the well-characterized erythroid-specific 3’HS1 CTCF element from the mouse β-globin locus (13), and showed that insertion of this site exhibited a prominent reduction in the level of YFP (Figure 2c and 2d). A similar effect was observed when we inserted the erythroid specific 5’HS4 CTCF element of the chicken β-globin locus (5) (Figure 2c and 2d), showing that the experimental system can quantify the activities of boundary elements even from other species.

We next tested CTCF binding elements from non-erythroid specific loci, including previously characterized CTCF elements located at the *H19-Igf2*, *HoxA*, and *Sox9-Kcnj2* loci (6,12,36,37), and a less prominent CTCF binding site located at the *Epha4-Pax3* TAD boundary (38). This showed that insertion of the CTCF elements from the *H19-Igf2*, *HoxA*, and *Sox9* loci strongly and significantly diminished the YFP level in the experimental system, while insertion of the less prominent CTCF site from the *Epha4-Pax3* TAD boundary reduced the YFP level to a lesser extent (Figure 2e and 2f).

Nearly all these diverse CTCF elements blocked the enhancer-promoter interaction and reduced alpha-globin expression to variable degrees and again all had a greater effect when the N-terminus of the bound CTCF protein was directed towards the enhancers (Figure 2). Together, these observations showed that the experimental system established here could quantify the activities of a wide variety of CTCF elements located on small fragments of 68bp including the proposed DNA contacts of CTCF and reflect their ability to block enhancer-promoter activities.

### Correlating the structure and function of individual CTCF elements

Previous studies of CTCF elements have correlated their activities with a variety of associated features including the sequence of the core CTCF binding element, its evolutionary conservation, the amount of CTCF and cohesin bound in ChIP assays, and their persistence following acute depletion of CTCF protein (18,20–24). While these parameters derived from large datasets may provide general trends, their predictive values have not been extensively tested on individual elements at the identical genomic position where all other variables are minimised.

First, by performing the FIMO analysis we found no correlation between the enhancer-promoter blocking activity of the CTCF elements and their resemblance to the consensus CTCF core motif described as position-specific scoring matrices (Figure 3a) (Spearman correlation, p=0.65). The degree of conservation of the CTCF element was also not correlated with their ability to alter enhancer-promoter activity as tested (Figure 3b) (linear regression, p=0.2171, R^2^=0.0935).

**Figure 3.**
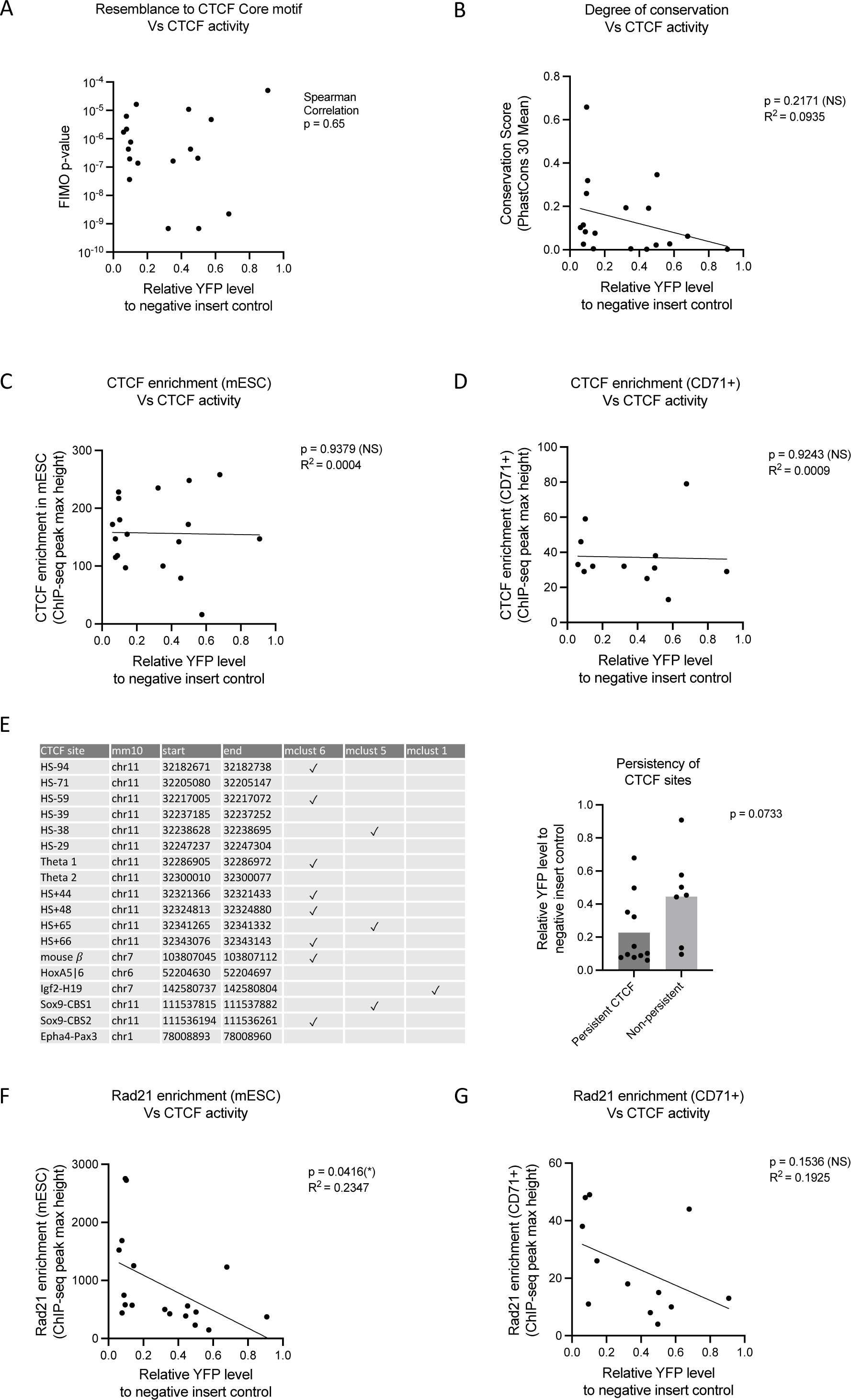
Correlating the structure and function of individual CTCF elements. (a) There was no correlation between the core CTCF binding sequences (p-value of the FIMO analysis) with their ability to alter enhancer promoter activity (Spearman correlation, p=0.65). (b) There was no correlation between the conservation of the inserted CTCF binding (PhastCons Vert30 genome conservation score) with their ability to block enhancer-promoter interaction (Linear regression, p=0.2171, R^2^=0.0935). (c) There was no significant correlation between CTCF activity and CTCF enrichment in the mESCs (linear regression, p=0.9379, R^2^=0.0004). (d) There was also no correlation between CTCF activity and CTCF enrichment when only focused on the erythroid-specific alpha-globin CTCF binding sites in the CD71+ erythroid cells isolated from EBs (linear regression, p=0.9243, R^2^=0.0009). (e) There was no association between the persistent CTCF elements and the enhancer-promoter blocking activity of individual CTCF sites (t-test, p=0.0733). Persistency groups is classified based on Luan et al. 2021 (24) (mclust 5 and mclust 6 are persistent CTCF sites; mclust 1 are non-persistent CTCF sites). (f) CTCF binding sites with stronger enhancer-promoter blocking activity showed a higher level of Rad21 enrichment in mESCs (linear regression, p=0.0416, R^2^=0.2347). (g) The correlation between the activities of the alpha-globin locus CTCF binding sites and enrichment of cohesin in CD71+ cells isolated from the differentiated EBs was not statistically significant (linear regression, p=0.1536, R^2^=0.1925).

Second, we investigated the relationship between the effect of each CTCF element and the levels of enrichment seen at these sites in their native context in the genome obtained from CTCF ChIP-seq data. In mESCs, we observed no significant correlation between activity and CTCF enrichment (Figure 3c) (linear regression, p=0.9379, R^2^=0.0004). Similarly, there was also no such correlation when we only focused on the erythroid-specific alpha-globin CTCF binding sites in the CD71+ erythroid cells isolated from EBs (Figure 3d) (linear regression, p=0.9243, R^2^=0.0009). There was also no correlation between so-called persistent CTCF elements (24) and the activity of individual elements when inserted between the enhancers and promoters (Figure 3e) (t-test, p=0.0733). These observations show that high levels of CTCF enrichment on CTCF binding sites at their normal genomic position do not necessarily predict their ability to block enhancer-promoter interaction.

Next, we interrogated the relationship between the enhancer-promoter blocking activity of CTCF elements and cohesin enrichment estimated from RAD21 ChIP-seq data. Interestingly, we observed that CTCF binding sites with stronger activity (i.e. lower relative YFP levels) showed a higher level of RAD21 enrichment in mESCs (Figure 3f) (linear regression, p=0.0416, R^2^=0.2347). As it has been shown that the loading and the enrichment level of cohesin may be tissue-specific (39,40), we have also correlated the activities of the alpha-globin locus CTCF binding sites and enrichment of cohesin in CD71+ cells isolated from the differentiated EBs. Although not statistically significant, we observed a trend that stronger CTCF elements (i.e. lower relative YFP level) tend to have higher levels of cohesin enrichment in their native loci (Figure 3g) (linear regression, p=0.1536, R^2^=0.1925). However, the levels of cohesin enrichment are likely to be affected by the position of each element in the native locus.

In summary, despite correlations found in genome-wide analyses, none of the commonly found associations fully predict the behaviour of individual CTCF elements in the sensitive assay used here to detect perturbations in the interactions between enhancers and promoters and the consequent changes in the levels of gene expression.

### Sequences flanking the core motif contribute to the strength of the CTCF element to perturb enhancer-promoter interactions

Although we found no predictive value for the CTCF blocking activity by analysing the core CTCF motif, it has been previously shown that sequences flanking the CTCF core motif may contribute to the functional role of a CTCF binding site. For instance, a phylogenetically conserved upstream motif has been discovered at 15% of all CTCF binding sites (20,41). The upstream motif was shown to stabilize CTCF occupancy by interacting with the zinc fingers of the CTCF protein at its C terminus (21,42). We scanned all the sites that we have tested and identified this upstream motif at the alpha-locus HS-94, HS-38, and HS+66 sites (Figure 4a). High resolution DNaseI foot printing we have published previously showed that the upstream regions of these sites are indeed bound by proteins in erythroid cells (26) (Figure 4b). Consistently, the sites associated with this upstream motif (HS-94, HS-38, and HS+66) exhibited very strong blocking activities compared to other CTCF binding sites analysed here (Figure 2a and 2b).

**Figure 4.**
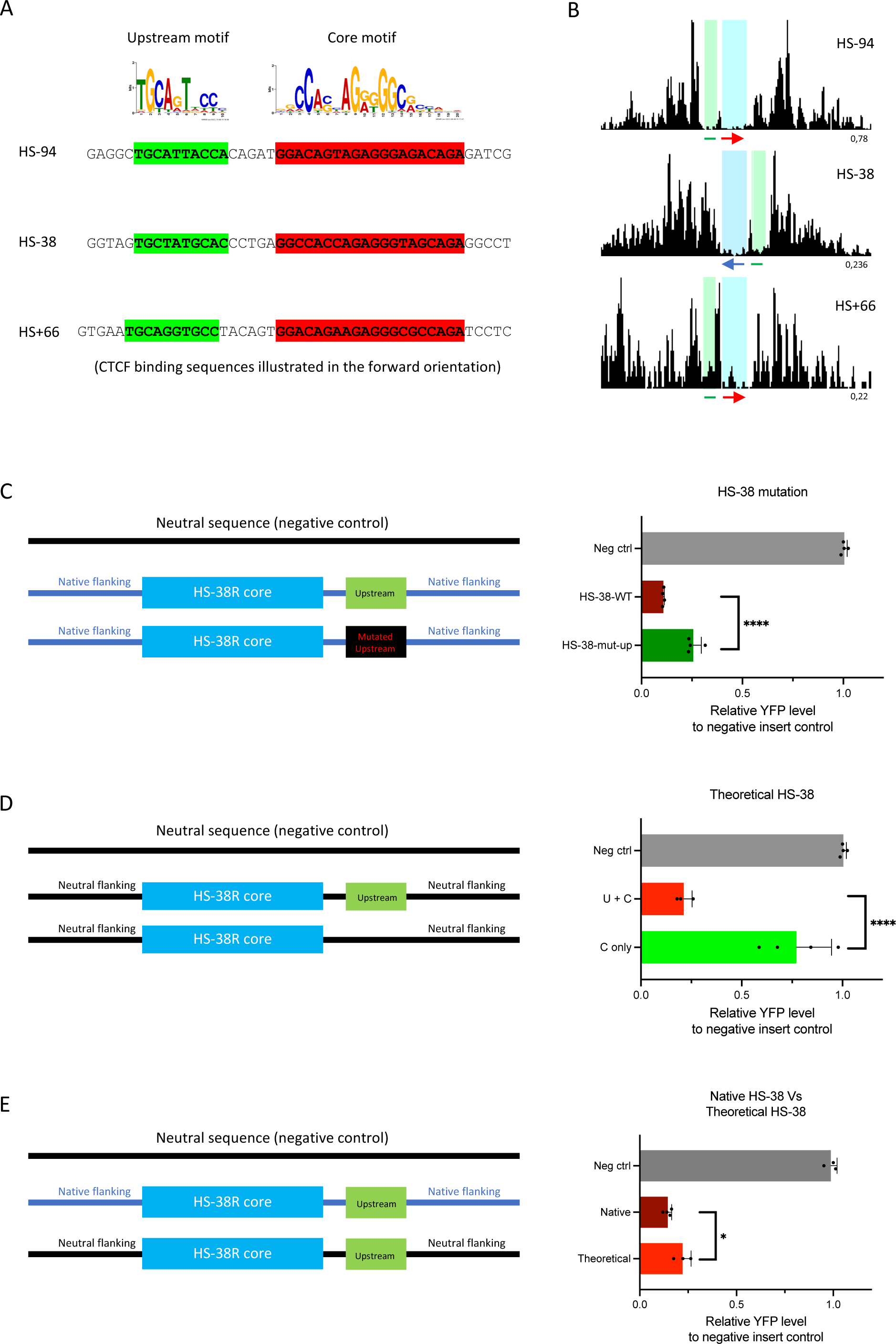
Sequences flanking the core motif contribute to the strength of the CTCF element in perturbing enhancer-promoter interactions. (a) The upstream and core motif sequences on the HS-94, HS-38, and HS+66 CTCF binding sites on the alpha-globin locus. (b) DNaseI footprinting data of the HS-94, HS-38, and HS+66 CTCF binding sites (Red arrow, forward oriented CTCF core motif; blue arrow, reverse oriented CTCF core motif; green line, known upstream motif; blue highlighted area, DNaseI footprints occupied by the CTCF core motifs; green highlighted area, DNaseI footprint occupied by the upstream motifs). (c) Mutation of the upstream motif of the native HS-38 site has significantly increased the relative YFP level when compared to the wildtype. (t-test, p<0.0001). (d) The relative YFP was remarkably increased when the upstream motif was absent in the theoretical HS-38 site. (t-test, p<0.0001). (e) The boundary strengths of the native and the theoretical HS-38 site are not equivalent. (t-test, p=0.0231).

To further explore the effect of the upstream motif on boundary strength, we synthesised artificial CTCF sites based on the alpha-globin HS-38 sequence and tested them in the CTCF activity reporter assay. First, we mutated the upstream motif of the native HS-38 sequence while keeping the core motif intact (Figure 4c). Mutation of the upstream motif significantly increased the relative level of YFP (indicating reduced insulator activity) when compared to the wildtype HS-38 site. Next, we investigated how the core and upstream motif sequences alone influence insulator strength without the influence of any native flanking sequences on HS-38. To achieve this, we synthesised theoretical CTCF sites by adding the HS-38 core motif plus upstream motif or the core motif only on a ‘neutral sequence’ from the alpha-globin locus. Again, the relative level of YFP was remarkably increased when the upstream motif was absent, indicating a significant reduction of the insulator strength (Figure 4d). These results show that the upstream motif sequence positively contributes to the insulator strength of CTCF binding sites.

Of interest, the newly synthesised HS-38 site did not fully recapitulate the boundary strength of the native HS-38 site. Even though the native and newly synthesised sites contain exactly the same core and upstream motif sequences, the boundary strength of the native HS-38 site is significantly stronger (Figure 4e) (p=0.023), suggesting that other sequences within the native site also contribute to the boundary activity.

To understand the underlying mechanism of the upstream motif, we have performed the CTCF ChIP-seq in the theoretical HS-38 models. Interestingly, we observed a drastic drop of the CTCF enrichment level in the absence of the upstream motif (Supp. figure 2). This suggests that the upstream motif sequences may strengthen the binding of the CTCF proteins on the element. It is also worth noting that the inserted theoretical HS-38 site containing the core and upstream motifs binds less CTCF proteins when compared with the native HS-38 site. This observation again suggests that the sequences flanking the core and upstream motifs in the native site could also contribute to the binding of CTCF proteins.

It therefore appears that at least one common motif immediately flanking the core CTCF motif in the small (64 bp) sequences studied here can modify the activity of the element. From our previously published DNaseI footprinting data, we have observed additional DNaseI footprints both upstream and downstream of certain CTCF sites such as the HS-59, HS+44, HS+48 and HS+65 on the alpha-globin locus (26) (Supp. figure 3). It remains to be seen if such sites may bind other proteins associated with insulator activity.

## DISCUSSION

To compare the ability of CTCF elements to insulate interactions between enhancers and promoters, we analysed uniformly sized sequences (68 base pairs) containing the core CTCF motif (20 bases) and 24 bases flanking upstream and downstream from 18 well-characterised elements. We show that regardless of their role in their natural genomic locations, nearly all CTCF elements analysed in this fixed, identical genomic location block enhancer-promoter interaction to different extents. In all cases the effect of the CTCF elements was greater when the N-terminus of the CTCF protein was orientated towards the enhancer.

Interestingly, the two *Sox9-Kcnj2* CTCF binding sites we have tested here have also been analysed using larger fragments (2.3-2.5 kb) in another study in which they were reported to have very subtle, or no boundary activities in a reporter assay based on the *Sox2* locus (28). Our observations suggest that the experimental system we have developed, based on the alpha-globin locus, may be more sensitive in quantifying the activities of CTCF-elements. Altogether, our 126 engineered models (encompassing the naturally occurring 18 CTCF sites, generated in both orientations, in addition to the mutants and artificial versions, each derived in three independent mESC clones) and the resulting observations demonstrate that the insulator assay based on the alpha-globin locus is a reliable and sensitive platform to quantify the activities of universal boundary elements in a non-biased manner.

We found no correlation between the ability of CTCF elements to block enhancer-promoter activity with the amount of CTCF or cohesin bound at the natural genomic sites of these elements; the degree of evolutionary conservation; or previously studied classes of CTCF elements based on the ranking of their core sequences (43). Nevertheless, using DNaseI footprinting we have shown that three of the strongest enhancer-promoter blockers include a previously described bound element lying upstream of the core CTCF binding sequence (20,41). In addition, we found other uncharacterised footprints located close to the core sequence that may affect function. This suggests that other proteins binding near to CTCF sites may contribute to insulator activity.

To maximize the testing capacity of the boundary reporter assay, we have recently miniaturized the differentiation of EBs to a 96-well format (29). Individual clones established in this study were cultured and differentiated in a 96-well plate which provides results consistent with the standardised model presented here (see supplementary appendix and supp. figure 4). This 96-well EB differentiation format will greatly improve the capacity of the experimental assay for higher throughput screening in future studies.

Together these findings show that, as with enhancers and promoters, using the approach outlined here, it will ultimately be possible to sub-classify CTCF elements in an unbiased way by sequence analysis independently of their roles in their natural genomic environment.

## DATA AVAILABILITY

ChIP-sequencing data generated for this study have been deposited in the Gene Expression Omnibus (GEO) under accession code GSE240678. Previously published ChIP-seq, DNaseI-seq and ATAC-seq data reanalysed here are available under the following accession codes: GSE97871, GSE30203.

All other data supporting the findings of this study are available from the corresponding author on reasonable request.

## SUPPLEMENTARY DATA

Supplementary table 1: Sequences of the homology arms of the CTCF element insertion site.

Supplementary figure 1: Establishing an experimental system to evaluate the capacity of individual CTCF elements to block enhancer-promoter interactions. (a) CTCF sites of interest were designed and synthesized. The synthesized fragment was cloned into the pROSA-TV2 HDR plasmid by Golden Gate assembly. The cloned HDR plasmid and the sgRNA plasmids that targets the landing pad between the enhancers and promoter of the alpha-globin gene were co-transfected into the YFP-tagged E14 reporter mES cells. The transfected cells were first subjected to Puromycin selection for the uptake of the sgRNA plasmids. The cells were then subjected to Hygromycin selection for the successful integration of the HDR fragment. The integrated hygromycin selection cassette was subsequently removed by Cre-LoxP excision. Finally, single colonies were isolated, and the correct clones were screened by the PCR-based approach. (b) A schematic diagram showing the position of the PCR screening primer pairs used to amplify the insert (1), or spanning the left (2) and right homology arms (3) to ensure the targeting occurred at the intended position at the alpha-globin locus.

Supplementary figure 2. Sequences flanking the core motif strengthen the binding of CTCF protein on the boundary element. CTCF ChIP-seq was performed in the theoretical HS-38 models. CTCF enrichment level was drastically decreased in the absence of the upstream motif (light blue highlighted region is the insertion site of the theoretical CTCF sequences). In addition, less CTCF proteins were enriched at the inserted theoretical HS-38 site containing the core and upstream motifs when compared with the native HS-38 site (red ChIP-seq track).

Supplementary figure 3. DNaseI footprinting data of the HS-59, HS+44, HS+48 and HS+65 on the alpha-globin locus shows additional DNaseI footprints both upstream and downstream of the CTCF core motif. (Red arrow, forward oriented CTCF core motif; Blue arrow, reverse oriented CTCF core motif; Grey line, unknown motifs underlying the additional footprints; blue highlighted area DNaseI footprints occupied by the CTCF core motifs)

Supplementary figure 4. EB differentiation can be scaled down to facilitate high-throughput CTCF activity measurement. (a) The trend of YFP level of individual CTCF insertion models yielded from the 10cm dish EB differentiation format. (b) The trend of YFP level of individual CTCF insertion models yielded from the 96-well EB differentiation format. (c) The YFP levels of individual CTCF insertion models yielded from the 10cm dish EB differentiation format correlated significantly with those yielded from the 96-well format. (linear regression, p=0.0002, R^2^=0.7289)

## ACKNOWLEDGEMENTS

We are very grateful to Lars Hanssen for the initial idea of the boundary element screening assay; Helena Francis, Andrew King and Danuta Jeziorksa for establishing the YFP-tagged hemizygous mESCs cell line; Lance Hentges for his advice on using LanceOtron; Prof. Jim Hughes, Prof. James Davies, Dr. Robert Beagrie and all the members of the Higgs lab for helpful comments on the manuscript; the Flow Cytometry Facility at the Weatherall Institute for Molecular Medicine (WIMM) for helping with the FACS sorting experiments.

