## Supplementary table 1 for "The characteristics of CTCF binding sequences contribute to enhancer blocking activity"

Supplementary Table 1. Sequences of the homology arms of the CTCF element insertion site.

| Left homology arms of the inserted site contained in the pROSA-TV2 vector: |
|---|
| CCTTGGCTATATATTGCACTTGAGGCTAACTCTAGATACACAAAACCCTGTCTGCCCAAAAATCA<br>AAGAAAAGGGAATAAATAAAAAATAACATAAGCAGCAAGCCCTTGAAGTGCAAATCTGCACGT<br>ACCGGCAACCCTGGGACACACACTATGTAGCTCAGCCTGACTGCAAACTCACAACAATCCTCCT<br>GCTTCAACCTCCACAACTGGAAGGTATTACAATCATATTCCACTACTCTTTAGTTTAAGATTACAA<br>ACCTGCCAAAAAGTGGTGGCACATACCTTTAATCCCAGCACTAGGGAGGCAGAGGCAGGCAGA<br>TCTCTGTGAATTTGAGGCTAGTCTCATCTACAGGGCTACTAGTTTCAGGATAGTCAGGACTACAC<br>AGCGAATCCTTGTCTCTCAAAAAAAAAAAAAAAAAAAAAGCAAACTGTTCACCACTATTCATCAG<br>AATAAGGCAGAATCGGAGCTATGACACTCTCAGTGTAGAGTGTCGGCTTCATACGCACAAGGCT<br>CAAGGTTTGGTTTCCAGTGCTAGGAGAGGCACTACACGCTACAGTGTTGGGGGGAGATAAAAA<br>TGTGTAAAAATGACAAAGAAATGATATCCATATTATTTTTCAAATTCAGACACATCAGTAAGACC<br>AGGACAACAGAAAAATAAGCATCTATAGAAGAAGAAGAAGAAGAAGAAGAAGAAGAAGAAG<br>AAGAAGAAGAAGAAGAAGAAGAAGAAGAAGAAGAAGAAGAAGAAAAAGAAGAAGAGG<br>AGGAGGAGAAAGAGAAAGAGAGAGAGAGAAAGAGAAAGAGAAAAAGAAAAAGAAAAAGAA<br>TCATCTAGATACATCAGGAAGAGGAAATCCAGTAGCCAGTGAAGTCATGAAAAAGGGCACAGC<br>CTCACTTACCACCAGGAAAGGAAATGGAAACCACAAGGGAATCACTGTGTCCTGACCAGAAAG<br>ACTAAATCAAAGGACACTAAGACCAGCTGTGCCAGATGTTAGGGAGGGAAACTCACACCAGTC<br>GCAGAAATGCAAATTGGCCACCCTTTGGAAAAGTCCCCCTTTCTTTTTCTAATGTTGAAAGTGCA<br>TGTGTCAGGACTGGCACATGGCTCGGGTGGGAACGTGCTTGCCTAGCCTGCAGGAAGCCCTGG<br>CTTCAGTCCCCAGCACCACGCAAACTGATTGTGGTGGCACATGCCTGCAATCTCAACCCTCAGG<br>AATTAGAGGCAGAAGGAGTACAAGTCCAGGGTCATCCATGGCTACCTAGGAAGTCTGAGGCTA<br>GCCTGGGATACATGAGACTCTGTCAGAGTTTGTGGTTGGGAGAGGAGGGAGGGAGAAGAAAG<br>GACACTTGGGTTGTTTCTACTTCAGGGTGTGATGGATAATTTTGTTGCAATGTCCCTGAATGAAG<br>G |
| Right homology arms of the inserted site contained in the pROSA-TV2 vector: |
| AACAATTAAAAGAGAATAACCGCTGTATTTGAAAGAAACAACTACAGTGATGCTCTTGCTATGC<br>ACACACTCCCATGTTTCATTTCTCCTGAGTGTATATGCAGTAATGGAAATAAAACAAGACCATAT<br>GGTAAGACCACCTACAATTTTTACCTATCAACCTGGCTGGCTGGCTGTCCATTGTCTATTTGTTTG<br>TTTGTTTATGGACAGGAGCTTTCTACATAGGATAGCCCTGGCTGTCCTGGAACTGCCTACACAGA<br>CCTGGCTGGTCTTAAGCTAAACTAAGATCTTCCTGTCTCTGCCTCCCAAGTGCTGGGATGGAAAG<br>TGTGCACCACCGTACCCTGCACCCTGCCTTTAACTTTTTGAGGAACTACCAAACTTCTTCCCCGT<br>GTGCACCCCTTTCTTTCTCTCCATTGCATCTTATACAGATTGCAATTTCTTTACATATTCTCTAGCA<br>TTTTTATTTTGTCTATTTGTTTGATTGTTTGTTTGTTTGTTTTTCAATACAGGGTTTCTCTGTGTAGCT<br>CTGGCTGTCTTGGAACTCATTCTGCAGACCAGTCTGGCCTGGAATTTCTCAGAGATCTGCCTACC<br>GCTGCCTCCCAAGTGTTGGGACTAAAGTCTTGTGCCACCATCCTGGCTTTATCTTGTCGTTTTAAT<br>TATGGCTATCCTAGTAGCTGTGAAGTAGTATATACAGCATTAAAAAAATTTTTTTAAAAGATTTAT<br>ATTTGTGAGTACACCGTTGTTGTCTTCAGACACACCAGAAGAGGAAATCAGATCCCATTACAAAT<br>GGTTGTAAGCCATCATGTGGTTGCTGGGAATTGAACTCAGAACCTCTAGAAGAGCAGCCAGTGC<br>TCTTAACCACTGAGCCATCTTTCCAGCTCCACATATACAGCATTTGTCTGTTTGTTTTTGATACAGT<br>GTCTCAACACTGTAACTTAGGTTAGCTTGCAGAAGTACTATCCAGTGTGTGACAATCAACATGAC<br>TTCCCTTCCTAAAGGCTGGGATCACAGACAGTCATCATCTTGTCTGGTGTTTATTTCCTTTGGAGA<br>AATAGTCAAAAATCCACTAGCTAATAAAAAGACAAACAGCTGTACATTCACATAGACAAATTTA<br>AAAAGAGACACTGGAGAAATGGCTCAGCGGTTAAAAGCACATGCTAATCTTCCAGAGGATCCA<br>AGTTTGGTTTTCTAGCACTCATATTAGCTGACTCACCTGTAACTCTGAACTCCAAGGGAT |
