## Supplementary figures and images for "The characteristics of CTCF binding sequences contribute to enhancer blocking activity"

### Supplementary figures

Supplementary Figure 1

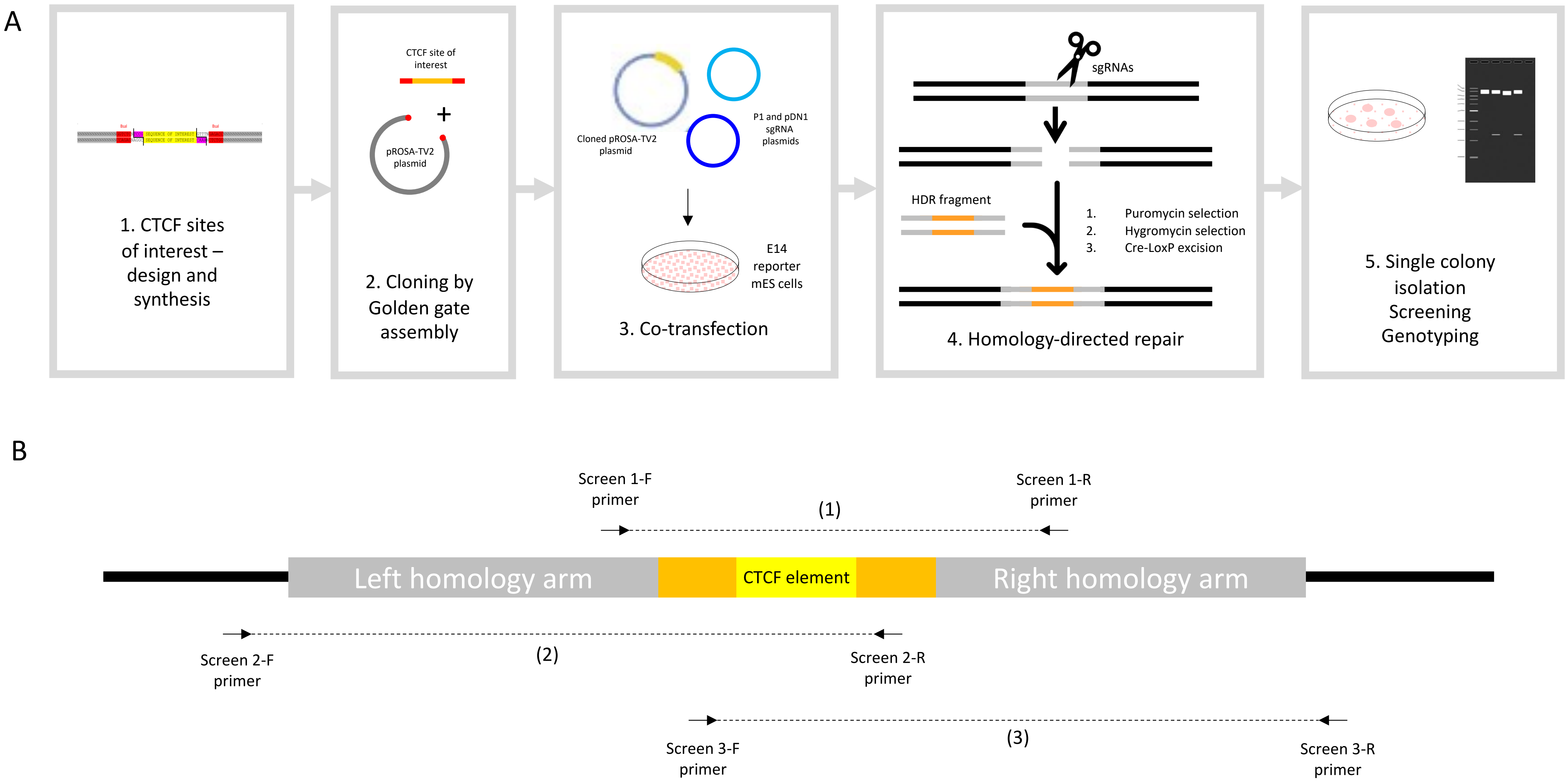

#### Supplementary Figure 2

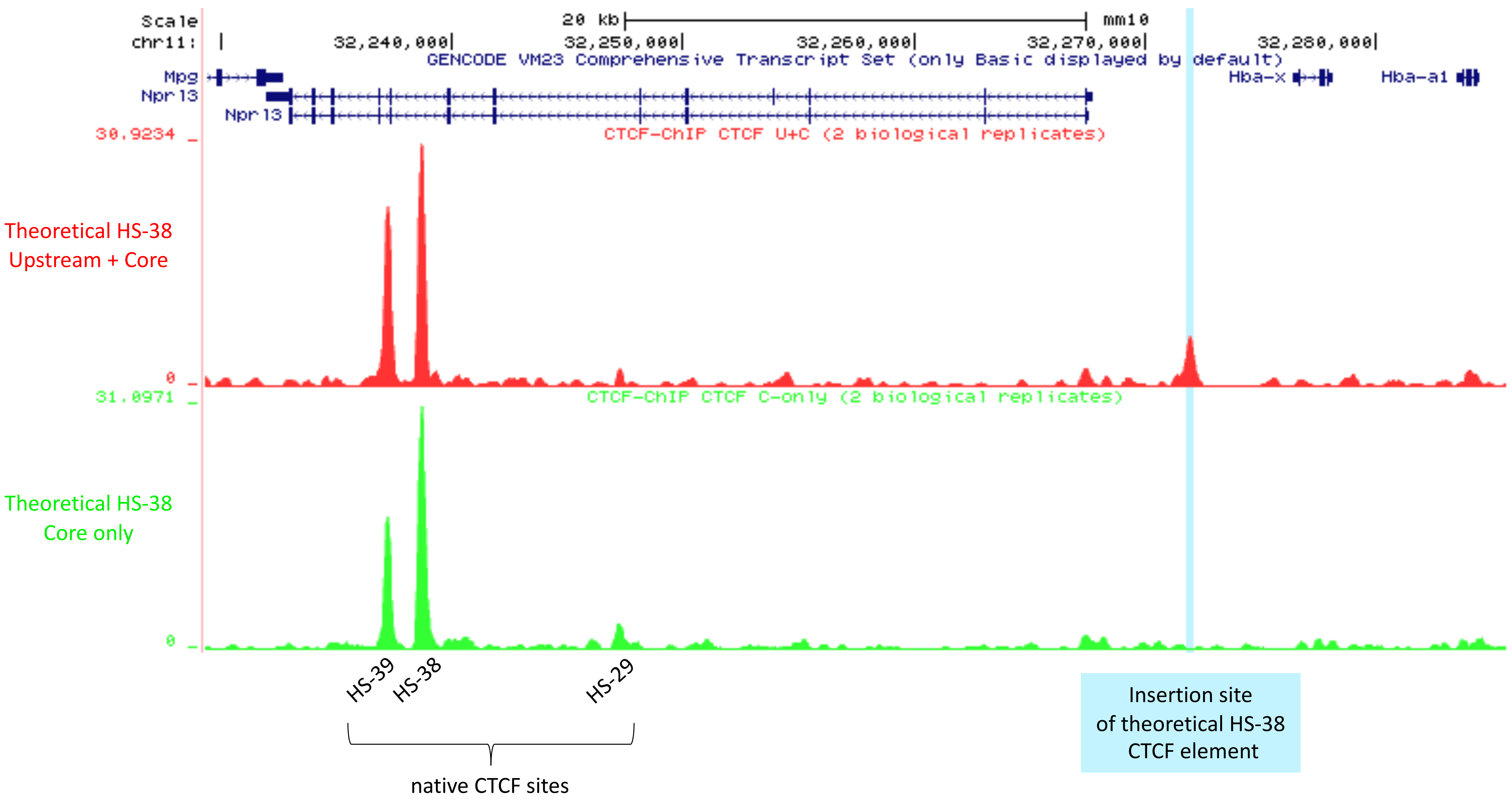

Supplementary Figure 3

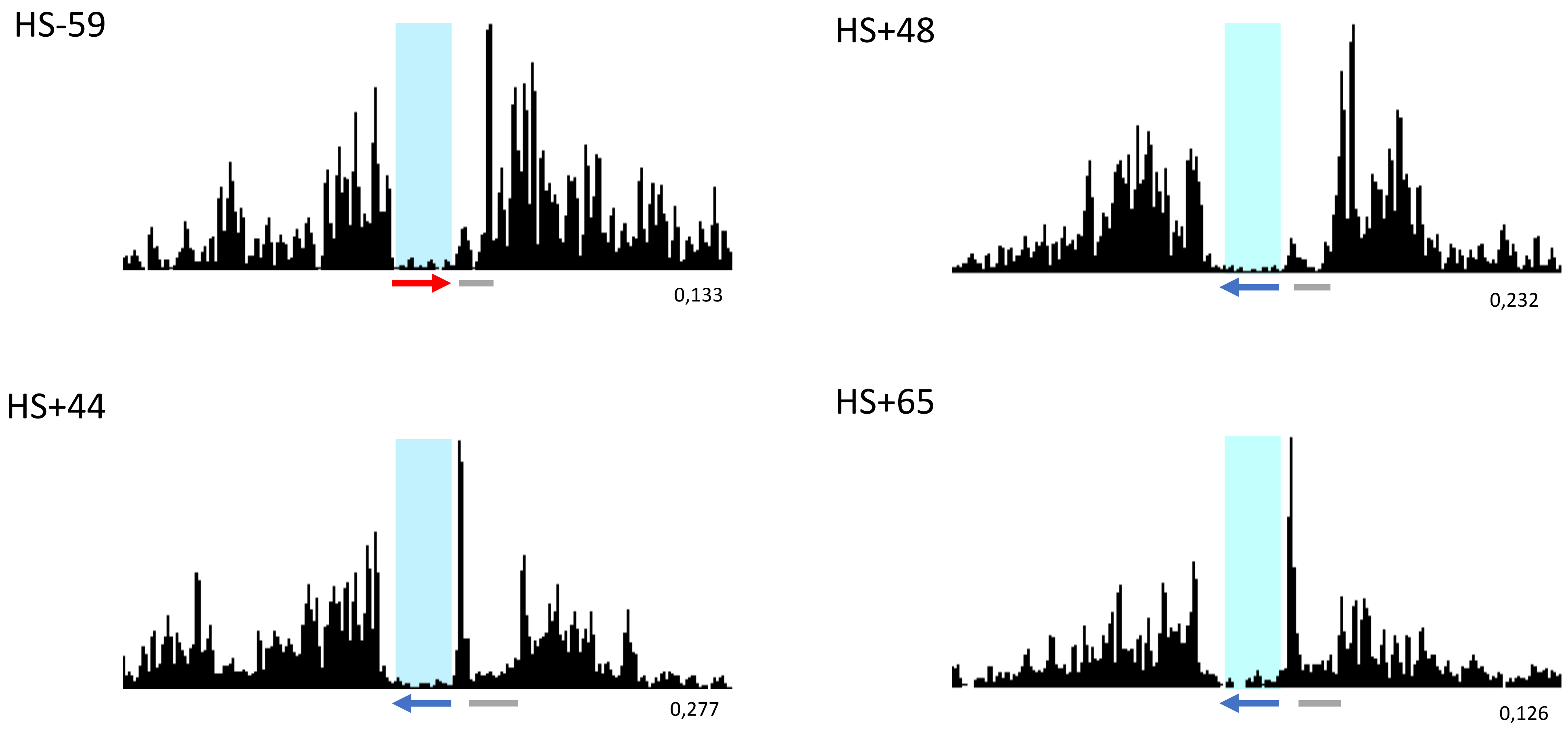

Supplementary Figure 4

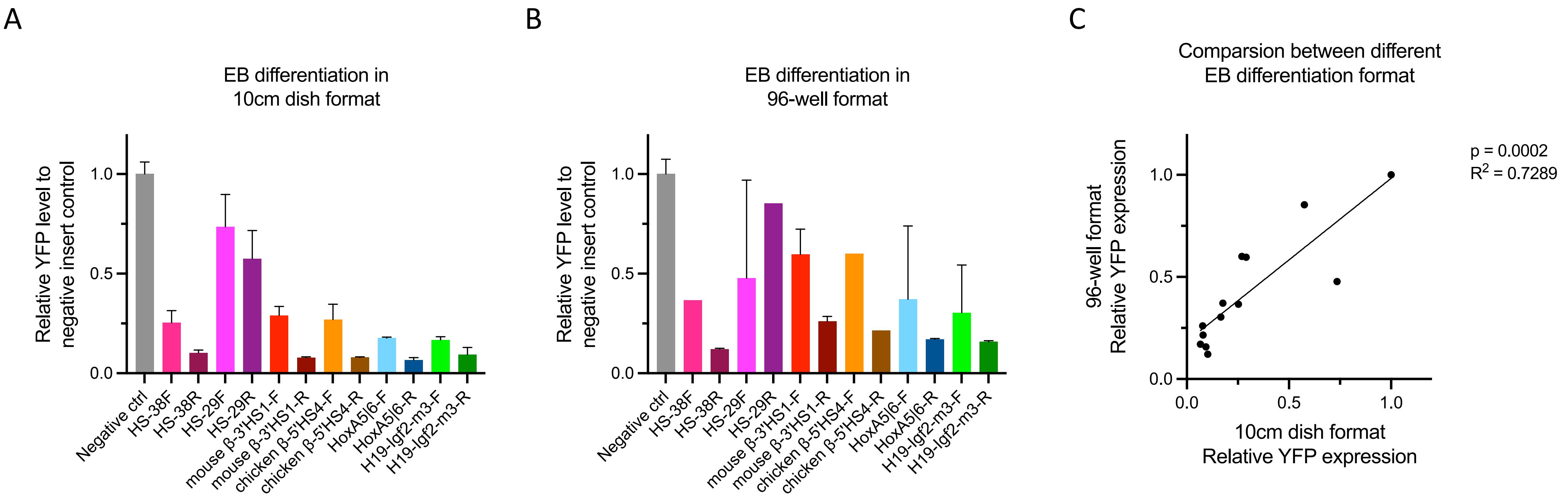
